# A dividing cell model for stable propagation and curing of *bona fide* human sporadic Creutzfeldt-Jakob Disease prions

**DOI:** 10.1101/2025.11.23.690013

**Authors:** Akin Nihat, Parineeta Arora, Christian Schmidt, Melissa L.D. Rayner, Jacqueline Linehan, Sebastian Brandner, Simon Mead, John Collinge, Parmjit S. Jat

## Abstract

Prions are assemblies of misfolded proteins that cause transmissible, progressive neurodegenerative disease in humans and other mammals. Human prion diseases have particular public health and biological significance, both due to their transmissibility between hosts, and the mechanisms they share with many other neurodegenerative conditions. However, their study has been considerably impeded by the absence of a cell-based system capable of reproducibly propagating and quantifying *bona fide* human prion infectivity, which has been a goal for several decades.

Here, we describe the first robust, cell-based assay of human prion infectivity in dividing cells. We found that CAD5 cells expressing human prion protein instead of the endogenous mouse prion protein, replicate human prion infectivity from brain samples of patients with sporadic Creutzfeldt-Jakob disease (sCJD) expressing valine at codon 129 of the prion protein gene. Moreover, these cells quantify human prion infectivity with similar sensitivity to the current gold standard animal bioassay, at a fraction of the cost and time, maintain persistent infection that can be cured with an anti-prion treatment, and can be adapted for therapeutic screening. This novel system permits direct cell-based quantification and investigation of human prion infectivity for the first time.

## Introduction

Prion diseases are a group of uniformly fatal, progressive neurodegenerative conditions caused by the autocatalytic, self-templated misfolding of the constitutive cellular prion protein (PrP^C^) into prions - assemblies of disease-associated isoforms, some of which acquire protease resistance (PrP^Sc^)^1^. Prion diseases occur naturally in humans and other mammals, including sheep (scrapie), cattle (Bovine Spongiform Encephalopathy, BSE) and cervids (Chronic Wasting Disease, CWD); they can also be inherited via mutations in the prion protein gene (*PRNP*), and acquired through environmental exposure (iatrogenic or experimental transmission, or ingesting infected material)^2^.

The most common form of human prion disease, sporadic Creutzfeldt-Jakob Disease (sCJD), has an annual worldwide incidence of 1-2 cases per million, increasing in recent decades^3^. Prion diseases remain of great public interest - both due to their zoonotic potential, and the increasing evidence of shared mechanisms between prion diseases and other neurodegenerative proteinopathies, such as Alzheimer’s disease^2, 4^. Prions exist as multiple strains, which cause discrete clinicopathological profiles that can be serially propagated, and determine host permissiveness to infection; these strains are composed of “clouds” of PrP protein conformational states^5^ that may be selected for under different environmental or genetic pressures^1^. A number of strains are recognised in sCJD, defined in part by the polymorphism at codon 129 (methionine/valine) of *PRNP*^6^.

Human prions are notoriously challenging to study, primarily due to the absence of a permissive cell system capable of reproducibly propagating them^7^. This has been mitigated in part by excellent transgenic mouse models expressing only human PrP^C^^8^, which faithfully replicate disease-related clinicopathological features. These models represent gold standard assays to determine infectious prion titre, probe human prion transmission and biology, and evaluate putative therapeutics, but are expensive, require significant numbers of mice and take months or years to deliver results.

Investigating prion pathobiology has therefore typically involved a combination of transgenic models, cell-free seeding assays such as Real-Time Quaking-Induced Conversion (RTQuIC) and Protein Misfolding Cyclic Amplification (PMCA)^9^, and cell systems that propagate non-human prions. Immortalised cells capable of maintaining infection with murine prions were described even before the prion pathogen itself was identified^10^. They provide relatively cheap, robust and reproducible systems to investigate prion biology, and crucially, *de novo* prion production can be differentiated from the inoculating material, which is diluted below the detection limit upon serial cell passage.

There now exist a plethora of dividing cell lines capable of propagating prions from a variety of non-human strains. Isolation of PK1 cells, a highly susceptible derivative of mouse N2a neuroblastoma cells, enabled the development of the Scrapie Cell Assay (SCA)^11^, which can be semi-automated for rapidly and precisely quantifying murine prions in medium to high-throughput. The SCA has formed the basis for key studies on prion propagation^12^, toxicity^13^ and structure^14^.

Whilst several fundamental observations regarding prion biology have emerged from the study of non-human prions, the absence of similar robust cell systems capable of propagating and quantifying human prions has proven to be a significant stumbling block. Anti-prion agents identified using high-throughput screening with non-human prions have failed to translate to efficacy in animal models or human trials^7^. Extensive evidence suggests that the mechanism of infection, cellular processing and clearance, and response to many anti-prion agents, can be strain-specific^15^ – identifying therapeutic compounds effective against human prions is therefore likely to require high-throughput systems able to directly propagate and quantify them.

Moreover, current international guidance regarding infectivity and transmission risk of human prions is based on low-sensitivity immunohistochemical analysis of post-mortem tissue^16^, yet recent evidence suggests the infectivity of several non-neuronal tissues is considerably higher, and more variable, than previously believed^17^. A systematic investigation of human prion tissue infectivity is all but impossible given the enormous number of transgenic mice required, in the absence of an appropriate cell-based assay.

The ability to persistently propagate, quantify and study human prions in a dividing cell-based paradigm has therefore remained a crucial goal for several decades after it was first outlined. CAD5 cells are a prion-permissive sub-clone of murine Cath.a-differentiated (CAD) cells, a central nervous system catecholaminergic line with neuronal properties^18^. They are susceptible to a wide range of murine prion strains^19^, and this tropism has recently been extended to hamster^20^ and CWD prions^21^ by expressing homologous PrP^C^ in place of the endogenous mouse protein. Transgenic mice expressing human PrP with valine at codon 129 instead of murine PrP, are susceptible to all human prion strains, regardless of *PRNP* codon 129 polymorphism^22^. We hypothesised that recapitulating this by stably engineering CAD5 cells to express human PrP with valine at codon 129 instead of the endogenous mouse prion protein, followed by serially single cell cloning, would generate a human prion-susceptible immortalised cell line. Here, we describe the derivation of a CAD5 cell line expressing human PrP that is the first reproducible system capable of propagating and quantifying infectious human sCJD prions in cell culture.

## Results

### Developing EKV cells, a CAD5 HuPrPmssV129 clone susceptible to sporadic CJD prions

As the starting point for human PrP reconstitution, we used CAD5-KDB3 cells, in which the mouse PrP has been stably knocked down using an shRNA targeting the 3’ untranslated region^23^. CAD5-KDB3 (KDB3) cells express PrP RNA at <1% level of CAD5 WT cells, and when reconstituted with the full length open reading frame of mouse PrP, are susceptible to 22L, RML, ME7 and MRC2 prion strains respectively^23^.

CAD5-KDB3 cells were reconstituted by retroviral infection with full length human PrP with valine at amino acid 129 and the mouse signal peptide sequence replacing the human equivalent, as we found this led to increased human PrP expression. VSV-G pseudotyped recombinant ecotropic retroviruses prepared from two independent constructs (pLNCX2-HuPrPmssV129) were used to reconstitute CAD5-KDB3 cells, and 240 stably transduced single cell G418-resistant clones isolated for each construct. Clones were expanded into cell lines and challenged with two sCJD-infected frontal cortex homogenates, and PrP^Sc^ detected in a modified Scrapie Cell Assay (SCA) format. 58 initial clones were selected and reduced over serial assay to 4 candidate clones, based on the detection of PrP^Sc^ over at least one cell passage after dilution of signal from the residual inoculum. These clones were next challenged with two sCJD-infected frontal cortex homogenates (T3MV and T2MM strain types, London classification^6^) in the SCA and assayed in replicate wells to identify an optimal clone, which yielded PrP^Sc^ spots that were clearly identifiable and accumulated over serial passages (Fig. 1A). The optimal clone, 81F9, demonstrated reproducible PrP^Sc^ accumulation when challenged with a 3x10^-4^ dilution of T3MV sCJD frontal cortex (Fig. 1B). In contrast, when clones were challenged with multiple T2MM sCJD homogenates, PrP^Sc^ deposition was limited, heterogenous and not reliably quantifiable, so cell line development was continued using sCJD homogenates from patients heterozygous or homozygous for valine at codon 129 (Types 2 and 3, London classification^6^, equivalent to types 1 and 2 in the widely used international classification system^24^).

**Figure 1:**
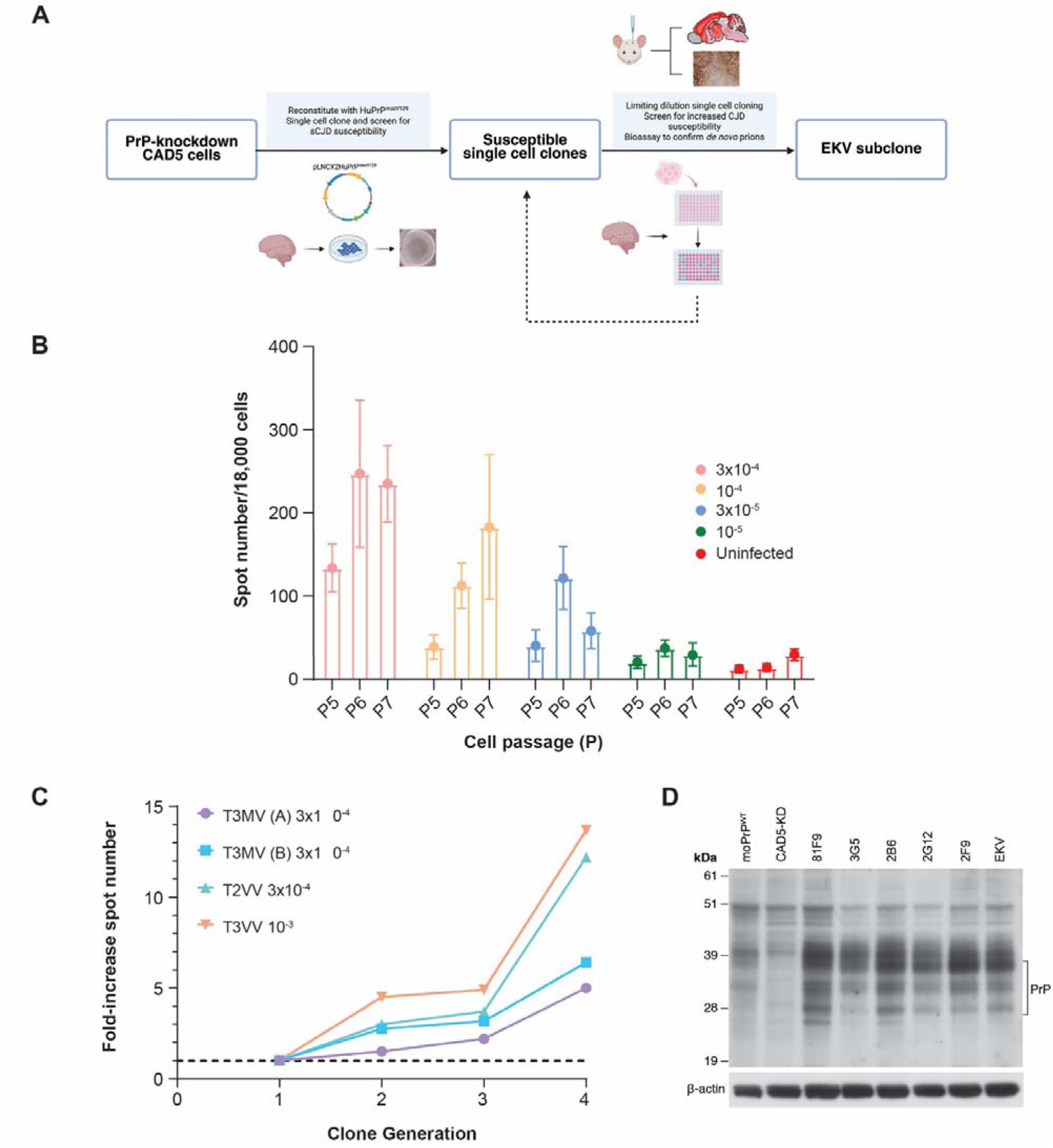
Iterative development of CAD5 HuPrPmssV129 clones susceptible to sporadic CJD prions. *A*, Schematic representation of the engineering, cloning and screening of CAD5 HuPrPmssV129 clones and development of the EKV cell line. CAD5-KDB3 cells, with mouse PrP expression knocked down, were reconstituted with the entire open reading frame of human *PRNP* (valine at codon 129) and the mouse signal peptide. Single cell clones were prepared and challenged with sporadic Creutzfeldt-Jakob (sCJD) T3MV/T2MM prions in an SCA format to identify susceptible clones. The SCA was repeated in a multi-well format to ensure the clones were reproducibly susceptible. One such clone, 81F9, was taken forward and iteratively single cell cloned and screened for increasing sCJD prion susceptibility, leading to EKV, the optimal clone. In parallel, sCJD-exposed 81F9 cells were subjected to Tg152c mouse bioassay to confirm propagation of *de novo* infectious prions. *B*, Susceptibility of the selected first-generation single cell clone, 81F9, to T3MV sCJD. Twelve replicate wells in 96-well format were challenged with four dilutions of sCJD T3MV frontal cortex homogenate (I4960, weight/volume dilutions), passaged seven times to clear the initial inoculum, and subjected to PK-digestion and Elispot revelation with ICSM18. Cell susceptibility was measured as PrP^Sc^ spot number per well and is displayed as mean spot number and standard deviation. *C*, Relative increase in susceptibility to infection with sCJD prions of the best clone from each generation of single cell clones. Four cell clones were challenged with T2/T3 sCJD prion-infected frontal cortex homogenate (valine hetero- or homozygous at codon 129) at the dilutions shown, passaged four times and subjected to Elispot assay. Mean spot count of 8 replicate wells per clone was normalised to fold-change from the 1^st^ generation clone, 81F9 (dotted line). *D*, Representative immunoblots of PrP^C^ expression in engineered CAD5 lines and subclones: CAD5 PrP-knockdown cells (CAD5-KD, lane 2), reconstituted with full-length MoPrP (moPrP^WT^, lane 1), the first selected HuPrPmssV129 clone (81F9, lane 3) and selected single cell clones with increased susceptibility to sCJD (3G5, 2^nd^ generation; 2B6/2G12, 3^rd^ generation; 2F9/EKV, 4^th^ generation). 15µg protein in 15µl was loaded per lane, blots were developed with anti-PrP antibody ICSM35 and stripped/reprobed with mouse anti-[actin antibody as a loading control.

Three further rounds of serial single cell cloning by limiting dilution were undertaken, challenging approximately 400-1000 clones per each round with increasing dilutions of sCJD-infected frontal cortex homogenate. After the fourth round, the best clone, EKV, exhibited a five to fourteen-fold greater response to sCJD homogenates than the initial clone, 81F9 (Fig. 1C).

Prion protein expression is obligatory but not sufficient for susceptibility to prion infection^25^. Transgenic mouse models expressing PrP at several times endogenous levels are associated with reduced incubation periods^26^, but overexpression in N2a^27^ and PK1^28^ cells failed to increase prion susceptibility, or render resistant cells permissive to infection. We next asked whether the sequential increase in cell clone susceptibility to sCJD could be explained by increased PrP expression. Immunoblotting for PrP expression in cell clones with increasing susceptibility to sCJD infection showed three clear migrating bands consistent with di-, mono- and unglycosylated forms of PrP^C^, at a greater level than in knockdown CAD5 cells reconstituted with mouse PrP (Fig. 1D, lane 1). However, consistent with previous findings, there was no clear correlation between increased susceptibility and greater PrP expression (Fig. 1D).

### Engineered CAD5 cells propagate *de novo*, infectious human sCJD prions

Infectivity is a defining feature of prions^29^, classically measured via the presence of proteinase K (PK)-resistant PrP^Sc^ aggregates in the Scrapie Cell Assay. It is clear that PK-resistant forms do not represent the totality of disease-associated prion species, many of which are variably protease-sensitive, and may also contribute to infectivity^30, 31^. Prion-susceptible cell lines can produce non-infectious, PK-resistant aggregates following PrP overexpression, or upon treatment with anti-prion agents^32^, so demonstrating production of *bona fide* prion infectivity in cell culture is crucial. Some cell systems determine prion propagation via seeded amplification assays such as RTQuIC^33^, which can measure seeding of non-infectious amyloid species, and may misrepresent infectivity^34–36^. We therefore used the gold standard animal bioassay to evaluate prion infectivity propagated in the engineered 81F9 cells.

81F9 and control CAD5-KDB3 (PrP-silenced) cells were infected with 0.03% sCJD T3MV frontal cortex homogenate (I4960), serially passaged six times (24 days post-treatment) and processed as per the SCA. 81F9 cells treated with sCJD (81F9 T+) cells accumulated PrP^Sc^ (mean well PrP^Sc^ spot number 247 ± 90), whilst treated CAD5-KDB3 (KDB3 T+) and untreated 81F9 (81F9 T-) cells did not accumulate PrP^Sc^ (mean spot number 7 ± 3 and 14 ± 5, respectively).

Pooled lysates from treated 81F9 T+, untreated 81F9 T-, and treated KDB3 T+ (PrP-null) cells were inoculated into groups of Tg(HuPrP129V^+/+^ *Prnp^0/0^*)-152 congenic mice. Tg152c mice express human PrP homozygous for valine at codon 129 at 4-8 times the level of normal human brain, on a mouse *Prnp*-null background^37^, and are susceptible to all human prion strains^22^. As a positive control, a separate group of Tg152c mice were inoculated with 1% I4960 sCJD T3MV frontal cortex homogenate (BH T+). Mice were monitored for clinical signs of prion disease and culled when these were identified, or at experimental endpoint (600 days post-inoculation, DPI).

All mice in the BH T+ and 81F9 T+ groups developed clinical signs of prion disease (Fig. 2A/B, BH T+: 9/9 mice, mean 235 days post-inoculation, SEM ± 15; 81F9 T+: 18/18 mice, mean 251 DPI, SEM ± 6). Immunohistochemical analysis for abnormal prion protein showed indistinguishable patterns between the two groups (Fig. 2C), demonstrating typical appearances of widespread moderate spongiosis (most pronounced in the thalamus and brainstem) and diffuse PrP^Sc^ synaptic deposition, with microplaques predominantly in the thalamus and midbrain. Molecular PrP^Sc^ typing of both groups following limited proteinase K (PK) digestion showed no difference in glycoform ratio or electrophoretic mobility (Fig. 2D).

**Figure 2:**
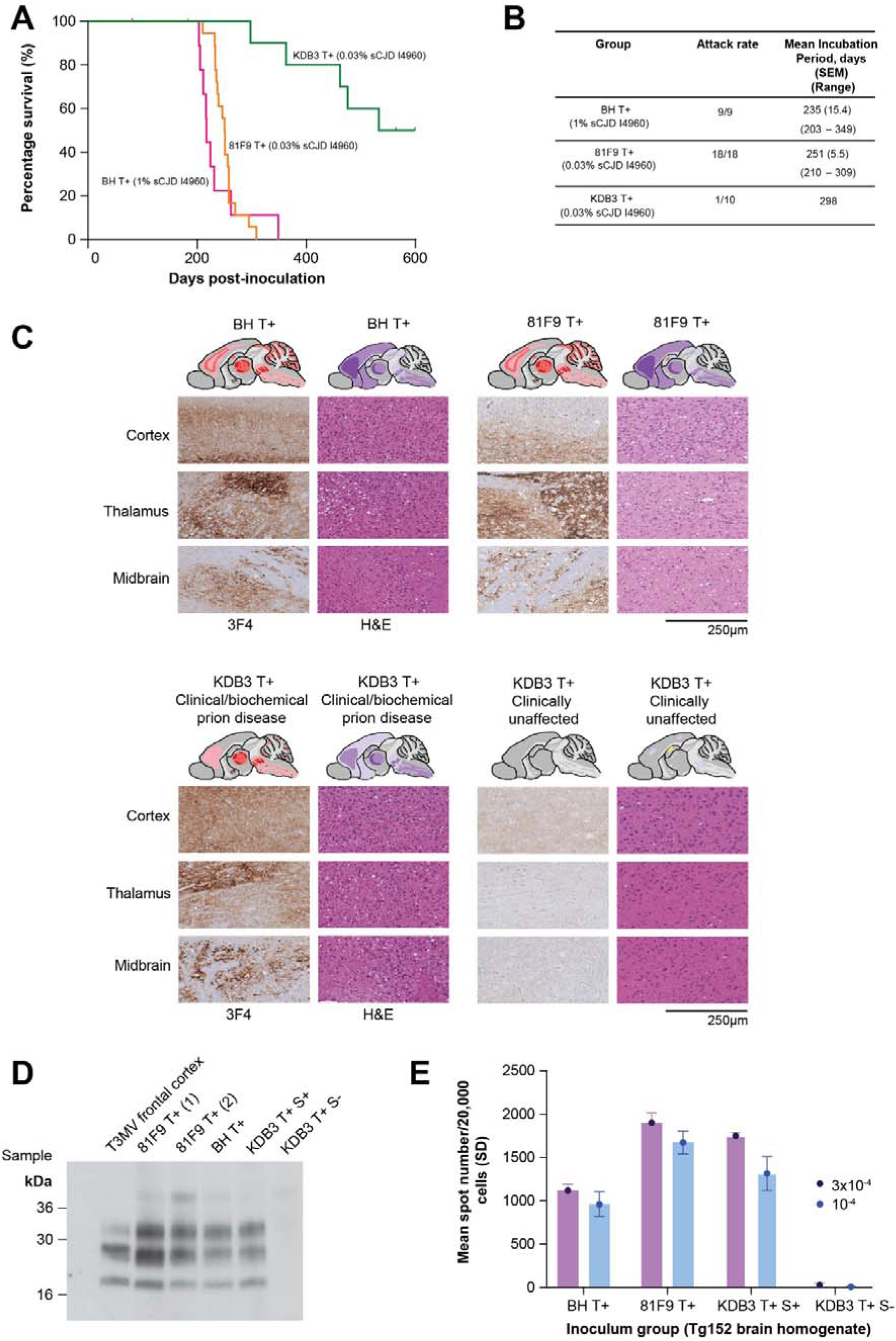
Engineered CAD5 cells propagate *de novo*, infectious human sCJD prions, with preserved biochemical properties. *A*, Kaplan-Meier survival curves for Tg152c mice inoculated with 1% brain homogenate from T3MV sCJD sample I4960 (BH T+), pooled 81F9 cell lysate after challenge with 0.03% I4960 homogenate and serial passage (81F9 T+), or CAD5 PrP^-/-^ KDB3 cell clone challenged with 0.03% I4960 homogenate (KDB3 T+) and serially passaged. Both BH T+ and 81F9 T+ curves were significantly different from KDB3 T+ (p<0.0001, log-rank with Bonferroni correction), but not from each other (p=0.30). *B*, Summary of clinical and histopathological attack rates, and mean incubation period by group. *C*, Neuropathological analysis of Tg152c (HuPrP V129) transgenic mouse brain after inoculation with sCJD-infected frontal cortex homogenate or sCJD-challenged cell homogenate. Each inoculum group is represented by three paired images of immunohistochemistry (anti-PrP antibody 3F4, left columns) demonstrating PrP deposition, and H&E stain (right columns) for neuronal loss and spongiosis, from cortical, thalamic and midbrain areas. Schematic drawings show the regional distribution of PrP deposition (diffuse synaptic staining in light red, darker shade indicates greater deposition; microplaques as dark red spots) and spongiosis (purple; mild-severe according to darker shade). Scale bar 250 µm; haematoxylin & eosin, H&E. *D*, Representative immunoblots of Tg152c brain homogenate from each group (81F9 T+ group represented with two samples, (1) and (2), from separate mice. I4960 T3MV sCJD frontal cortex shown in lane 1 as a control. *E*, Inoculation of Tg152c mouse brain homogenate into prion-naïve, highly-sensitive EKV cells to assess for occult infectivity; KDB3 T+ S+, brain homogenate from the single mouse with symptomatic and histopathological prion disease from the KDB3 T+ group; KDB3 T+ S-, representative brain homogenate from KBD3 T+ mice (n=9), without histopathological prion disease. Displayed are means/standard deviations of 6-well replicates from two independent experiments, brain homogenate challenge at two dilutions (3x10^-4^ and 10^-4^).

From the group inoculated with lysate from PrP-null KDB3 cells challenged with 0.03% I4960 (KDB3 T+), one mouse developed clinical signs suggestive of prion disease (at 298 DPI), with immunohistochemical evidence of prion disease (spongiosis and PrP^Sc^ deposition), and PK-resistant PrP^Sc^ on immunoblotting (Fig. 2A/B, KDB3 T+ panels). Other KDB3 T+ mice had no evidence of histopathological hallmarks of prion disease, nor PrP^Sc^ deposition (Fig. 2C and D, KDB3 T+ Clinically unaffected), although 4 other mice were culled due to unrelated health concerns after day 350 post-inoculation.

We considered that this group of KDB3 T+ mice culled due to health concerns could harbour occult prion infectivity, despite having no demonstrable PrP^Sc^ deposition or spongiosis. To test this, we exposed highly-susceptible, 4^th^ generation EKV cells to brain homogenate from mice culled from each group of the bioassay (Fig. 2E). As expected, treatment with homogenate from BH T+ or 81F9 T+ group mice resulted in stable accumulation of PrP^Sc^ spots after several cell passages, as did brain homogenate from the single PrP^Sc^-positive KDB3 T+ animal. None of the KDB3 T+, PrP^Sc^ and histopathology-negative brain homogenates induced PrP^Sc^ accumulation in EKV culture. In our extensive experience, Tg152c mice do not spontaneously generate prions. Together, this suggests that only 1/10 of the mice treated with KDB3 T+ pooled cell lysate became prion-infected, and that this cell lysate harboured a very low level of residual prion infectivity from the original brain homogenate after six cell passages.

### Establishing a cell-based Human Prion Assay (HPA) to measure sCJD infectivity from human brain

Having confirmed that 81F9 cells propagated *bona fide* infectious human prions, we next asked whether EKV, a more susceptible derivative of these cells, could be adapted to a reproducible cell-based assay of human prion infectivity (the Human Prion Assay, HPA) – analogous to the mouse Scrapie Cell Assay (SCA).

We optimised growth and prion challenge conditions for EKV cells (data not shown), and initially investigated the linear and dynamic range of PrP^Sc^ detection, using aliquots of a single stock of T3MV sCJD frontal cortex homogenate (I4960). EKV cells were challenged with serial 1:3 homogenate dilutions, in seven assays over a period of nine months (Fig. 3A). As expected, there was a degree of inter-assay variability in absolute PrP^Sc^ spot count, likely due to minor variations in homogenate, cell response, experimental reagents and other assay conditions^32^. However, across all assays there was a linear detection range of at least 1 log of inoculum concentration, whilst the dynamic range extended to 2 logs (10^-3^ to 10^-5^, or approximately 40 – 1000 PrP^Sc^ spots). The signal to noise ratio was excellent (<10) up to homogenate concentrations of 10^-5^ in all assays, corresponding to an ability to detect infectious prions from the sampled T3MV I4960 crude brain homogenate when diluted 100,000-fold (mean assay lower limit of detection 1.02^-5^ dilution).

**Figure 3:**
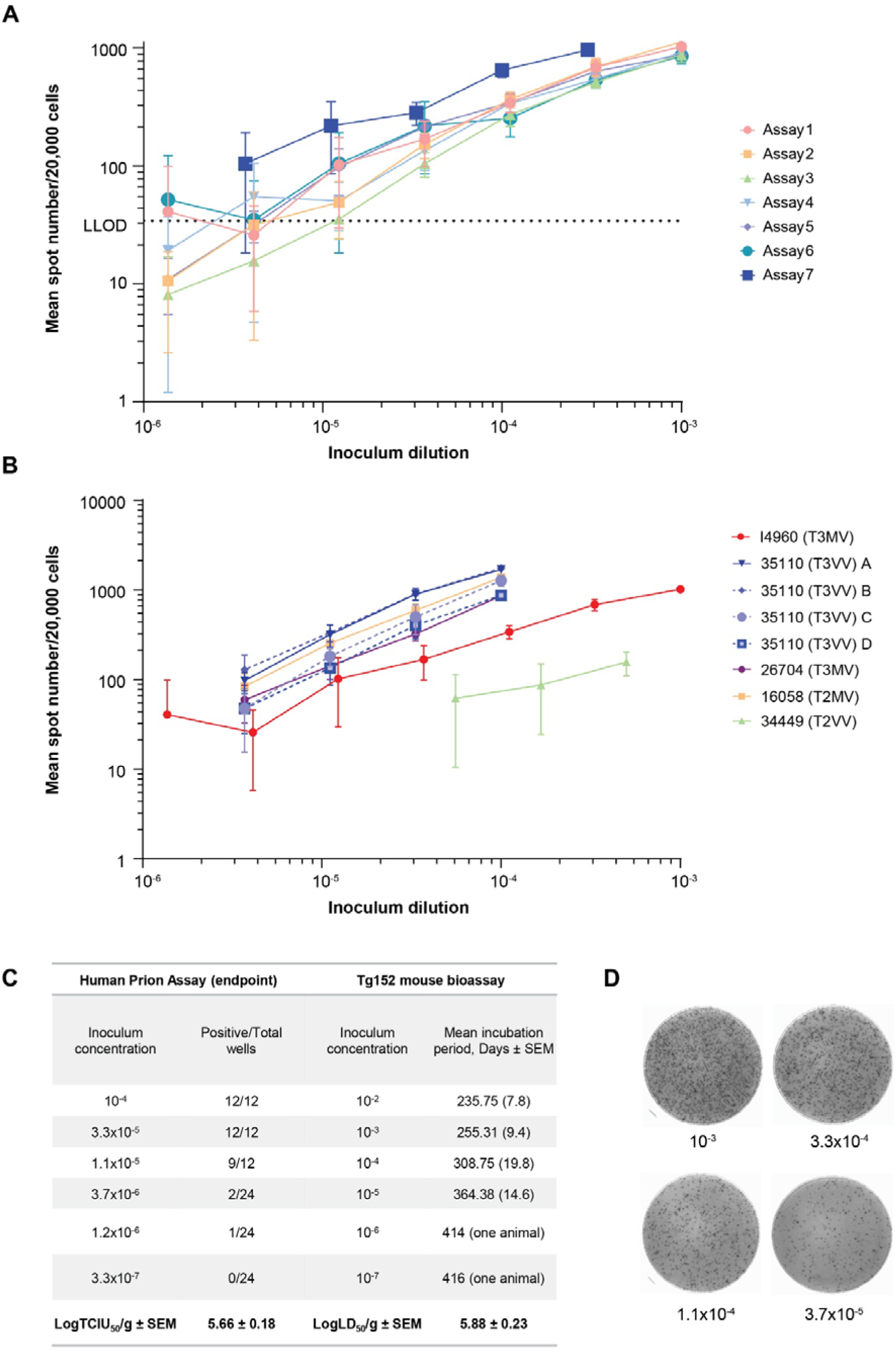
Establishment of a cell-based Human Prion Assay (HPA) to measure sCJD infectivity from human brain. *A*, Linear and dynamic range of the HPA over serial assays. Seven independent assays were undertaken over 9 months, using separate aliquots of the same sCJD frontal cortex homogenate (T3MV, I4960). EKV cells were challenged with serial 1:3 dilutions of I4960 (12 wells per dilution, 10^-3^ - 1.37x10^-6^), passaged 4 times and infectivity quantified (mean PrP^Sc^ spot number and standard deviation per 20,000 cells). HPA lower limit of detection (*LLOD, 34 spots per well*) was calculated as the pooled mean of uninfected wells (12 wells per assay, 20,000 cells per well filtered to Elispot) + 3 x pooled standard deviations. HPA linear range (3.3x10^-4^ to 1.1x 10^-5^) was defined as the inoculum dilution range resulting in a strong linear relationship (R^2^ > 0.9, mean across seven assays 0.94 ± 0.02) between PrP^Sc^ spot number and dilution above LLOD, across all assays on a double logarithmic plot. *B*, HPAs measuring infectivity from five different sCJD frontal cortex homogenates (T2/3, MV/VV). EKV cells (12 wells per dilution) were challenged with serial dilutions of five sCJD frontal cortex homogenates: four independent assays with different homogenate aliquots of 35110 (T3VV); I4960 (Assay 1 from Fig. 1a); 26704 (T3MV); 16058 (T2MV); 34449 (T2VV). Cells were passaged 4 times and PrP^Sc^ quantified as outlined previously. Dotted lines represent different aliquots of I4960 and 35110, assayed in independent experiments. *C*, Comparative sensitivity and estimated sCJD infectious titre by endpoint HPA and gold-standard transgenic mouse bioassay. In the endpoint HPA, EKV cells were challenged with 300 µl serial 1:3 dilutions of I4960 (T3MV) sCJD frontal cortex homogenate (12-24 wells per dilution), passaged seven times and PrP^Sc^ quantified; wells were recorded as positive if the spot number was greater than the mean spot number of uninfected EKV cells + 10 x standard deviations (12 wells per assay). Displayed are the number of positive and total wells per dilution from one of three independent experiments. For the mouse bioassay, Tg152c mice expressing human PrP (valine at codon 129) were inoculated intracerebrally with 30µl serial 1:10 dilutions of I4960 brain homogenate (three groups of 20 mice per dilution, each group inoculated with a different aliquot). The mean incubation period per group is shown for serial inoculations of one of three I4960 aliquots into groups of 20 mice. Infectious titres were calculated using a modified Karber method^64^, expressed as log tissue culture infectious units (LogTCID_50_/g brain homogenate) for the HPA (mean of three independent experiments ± SEM), and log lethal dose (LD_50_/g brain homogenate) for the mouse bioassay (mean of three inoculation groups, ± SEM). *D*, representative Elispot wells showing PrP^Sc^-positive spots for EKV cells challenged with serial dilutions of I4960 brain homogenate.

Next, we examined the response of EKV cells to challenge with sporadic CJD prions from a range of homogenates, strain types and codon 129 genotypes. EKV cells were challenged with serial dilutions of a panel of frontal cortex homogenates containing Type 2 or 3 prions^6^, codon 129 genotype MV or VV. Cells were susceptible to infection with all assayed MV/VV homogenates (Fig. 3B), with a demonstrable dose response that varied by up to 2 logs of concentration between homogenates. This was maintained over infection with three different aliquots of either I4960 (T3MV) or 35110 (T3VV) frontal cortex homogenate; whilst absolute spot count varied between assays, the relative count between homogenates remained consistent (R^2^>0.97 each assay) – suggesting persistent differences in infectivity between homogenates. EKV cells showed no clear PrP^Sc^ deposition in response to challenge with Type 2 sCJD-infected brain homogenate from patients with the MM codon 129 genotype (data not shown).

Similar to the SCA, the HPA can be performed in endpoint format to establish the infectious titre of samples (Tissue Culture Infectious Units required to infect 50% of microplate wells, TCIU_50_). Cells are challenged in replicate at limiting dilutions of brain homogenate - close to the endpoint dilution, only a small number of cells become infected, but over several generations PrP^Sc^ may become detectable as these cells may iteratively transfer infectivity to daughter cells provided they are not passaged too harshly^11^. Wells that ultimately surpass the non-infected background are deemed “positive”, and used to calculate TCIU_50_. We challenged EKV cells in 12-24 well replicates with serial 1:3 dilutions of I4960 brain homogenate, passaged cells seven times and determined the PrP^Sc^ spot count. Serial 1:10 dilutions of the same homogenate were inoculated into three groups of 20 Tg152c mice expressing human PrP with valine at codon 129, to establish a comparative gold standard infectious titre required to infect 50% of mice (ID_50_). Endpoint titration in EKV cells estimated the I4960 homogenate infectivity titre as 5.7 ± 0.2 logTCIU_50_/g, comparable with the mouse bioassay-derived titre of 5.9 ± 0.2 logLD_50_/g (Fig. 3C). However, whilst the mouse bioassay required significant sacrifice of animals and took approximately 500 days, the cell-based bioassay was over tenfold faster, without sacrificing animals, and at a fraction of the cost.

### EKV cells maintain a chronic human prion infection, which is cleared upon treatment with an anti-prion protein monoclonal antibody

We next sought to use EKV cells to develop cells that are persistently infected with human sCJD prions – as a platform for high throughput screens of therapeutic agents that can interfere with the propagation of infectious human prions.

We adopted a method used successfully to develop chronically-infected cells propagating non-human prions^38^. EKV cells were challenged with two sCJD homogenates, I4960 (T3MV) and 16058 (T2VV), passaged five times to clear residual inoculum, and then single cell cloned by limiting dilution such that on average one cell was seeded per well of a 96-well microplate. When wells are expanded to confluence, single infected cells may form a clonal population that persistently propagates infectious prions. We identified several infected EKV (iEKV) subclones with high PrP^Sc^ load, which were assayed over further passages in a dilution series to estimate persistence and proportion of PrP^Sc^-positive cells: clones 1G2 (T3MV, 50% ± 10% cells infected) and 1E10 (T2VV, 59% ± 14% cells infected) were selected for further assays. Both clones maintained stable prion propagation up to 15 cell passages (40-55 days, Fig. 4A and B), and could be cryopreserved and resuscitated without loss of prion infectivity.

**Figure 4:**
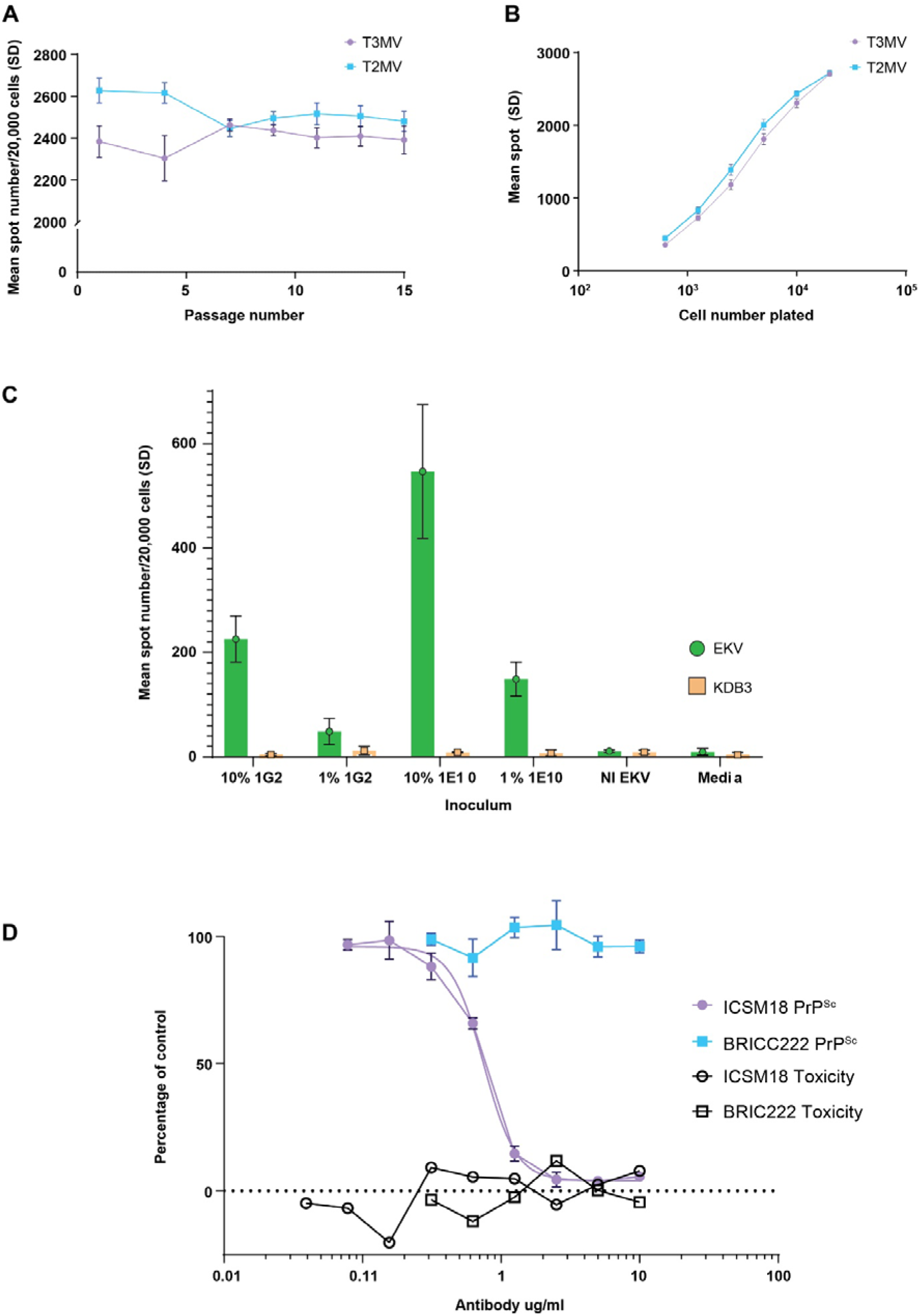
EKV cells can persistently propagate infectious human prions, and demonstrate efficacy of an anti-prion agent. *A*, Long-term propagation of human prions in EKV cells. Two selected iEKV subclones infected with T3MV (1G2) and T2MV (1E10) prion samples were serially passaged in 6-well replicates, and PrP^Sc^ assayed on alternate passages (20,000 cells). *B*, PrP^Sc^ spot count in 1G2 and 1E10 clones is a function of plated cell number. PrP^Sc^ load was quantified in serial 1:2 dilutions of 1G2 & 1E10 clones in 6-well replicates; mean PrP^Sc^ spot count (representing a single infected cell) for 1G2 was 50% ± 10% total cell count, and 59% ± 14% for clone 1E10. *C*, Cell homogenate from iEKV clones can transmit infectivity to naïve EKV cells. 10^7^ cells of each clone were resuspended in 1ml D-PBS and ribolysed. Uninfected EKV cells were challenged with 10% and 1% cell homogenates in media passaged four times and PrP^Sc^ quantified, as described. NI EKV, Non-infected EKV cells were controls. *D*, Treatment of 1G2 iEKV cells with anti-prion monoclonal antibody ICSM18. At confluence, 10,000 1G2 cells were seeded in 6 replicates to wells of a 96-well microplates, allowed to settle for 2 hours, then incubated with a dilution series of ICSM18 or its isotype control, BRIC222 antibody. After 5 days, media was gently replaced and 7,000 cells per well filtered for PrP^Sc^ ELISPOT detection, or cell viability assay. Untreated iEKV cells, non-infected EKV cells, media-only wells and DMSO-treated iEKV cells were used as experimental controls (not shown). PrP^Sc^ and toxicity were normalized as a percentage of untreated iEKV cells in 4-6-well replicates. Displayed are representative results from one of three independent experiments.

To confirm chronic production of infectious prions, we determined whether iEKV cell homogenates could infect naïve EKV cells. 1G2 and 1E10 cell suspensions were ribolysed and 1% and 10% homogenates used to infect EKV and PrP-null KDB3 cells in an HPA. Homogenates from both clones successfully infected EKV, but not PrP-silenced KDB3 cells (Fig. 4C), confirming the propagation of *bona fide* prion infectivity in both iEKV cell clones.

To support development of future therapeutic assays, we undertook a pilot treatment trial using the mouse anti-prion protein monoclonal antibody ICSM18, which has been shown to inhibit propagation of Rocky Mountain Laboratory (RML) prions in culture^39^ and *in vivo*^40^. 1G2 cells were treated for 5 days with serial 1:2 dilutions of ICSM18 (10 - 0.08 µg/ml) or BRIC222 (10 – 0.156 µg/ml), an IgG1 isotype control mouse monoclonal antibody, and PrP^Sc^ levels and cytotoxicity measured (Fig. 4D). ICSM18, but not BRIC222, depleted PrP^Sc^ in a dose-dependent manner. After 5 days of treatment with 10 µg/ml (67 nM) ICSM18, >90% PrP^Sc^ was cleared, without increased cell toxicity. Over three independent assays, the ICSM18 mean EC_50_ was 4.7 nM ± 0.40, which is approximately 6-fold higher than the EC_50_ reported in iPK1 neuroblastoma cells infected with the RML strain of mouse prions^39, 41^.

## Discussion

Developing an immortalised cell system capable of persistently propagating and quantifying infectious human prions has been a major goal of the prion field for several decades^7^. Here, we have established a dividing CAD5-derived cell line expressing human PrP^C^ that can be reproducibly infected with human sCJD prions expressing two codon 129 genotypes, remain stably infected upon extended culture and cryopreservation, quantify infectivity over two logs of concentration, and demonstrate the effect of an anti-prion treatment. Moreover, for the first time we show that prion propagation in cells produces *bona fide*, *de novo* human prions, which are infectious in a transgenic mouse bioassay, and in cell culture assays. This cell system has a wide range of potential uses in prion disease research.

The only previous report of human prion propagation in an immortalised cell line was unfortunately never replicated^42^. More recently, PrP^Sc^ accumulation has been described in terminally-differentiated cells exposed to human prions: human induced pluripotent stem cell (iPSC)-derived astrocytes produce PrP^Sc^ after challenge with sCJD (MM/VV) and variant CJD prions^43^, and human iPSC-derived cerebral organoids respond to sCJD MV brain homogenate^44^. These models are valuable tools to probe basic human prion biology, particularly as the latter systems provide a human cellular microenvironment. However, there are a number of limitations to their practical use. They are technically challenging to prepare and maintain, each experimental cycle requires several months and new prion-infected brain homogenate samples, and the cell cultures are heterogenous; these factors mitigate against their use for medium or high-throughput screens^45^. Additionally, because non-dividing cells cannot be serially passaged to dilute out residual inoculum, *de novo* infective prion production cannot be reliably differentiated or quantified.

We have shown that EKV cells can reproducibly measure infectivity in VV/MV sCJD prion-infected brain homogenates with similar sensitivity to the gold standard transgenic mice assay, offering an alternative that requires no use of animals, is considerably faster, and cheaper. Chronically-infected iEKV cells offer a persistent supply of human sCJD prions with preserved biochemical fidelity, and may reduce the necessity for human brain homogenate samples in future experiments. They can be effectively cured with an anti-prion protein monoclonal antibody; we are currently adapting the system to a semi-automated liquid-handling system that should allow high throughput screening of compounds that interfere with human prion replication for the first time.

The specific factors and conditions required for productive cellular prion infection remain frustratingly opaque. Homologous PrP^C^ expression is clearly necessary, but neither adequate nor correlated with degree of susceptibility^46, 47^; indeed, we found no consistent increased expression in more susceptible subclones. In immortalised cells, the vast majority of cell-cell transmission appears to occur vertically^48^, so a favourable balance between prion depletion through cell division and the intrinsic replication rate is axiomatic^25, 49^ – however, these conclusions were drawn exclusively through study of non-human prions. It has been argued that the replication rate of human prions may be too slow to permit propagation in dividing cells^50^. In this work, we did not observe a prominent difference in doubling time between the more or less prion-permissive subclones (data not shown), suggesting this was not a major driver of increasing susceptibility.

Previous investigation of immortalised, prion-permissive cell lines points to the importance of PrP^C^ availability at the cell surface^51^, and extracellular matrix composition^28^, but no unifying factors have been described^52^. It is noteworthy that cellular permissiveness to prion infection is highly strain-specific; different strains rely on diverse cellular pathways for uptake, infection, trafficking and replication rate^53, 54^. It will be valuable to determine whether human prions show such diversity in cellular processing in EKV cells.

Given their wide prion tropism, there may be CAD5-specific factors that are particularly supportive of prion infection /propagation. Expressing human PrP^C^ in RK13 cells, a highly prion-susceptible cell line with no endogenous PrP^C^ expression, failed to support human prion propagation^55^; as did humanised N2a cells, which are susceptible to murine prions when expressing cognate PrP^C^ (PA and PSJ, unpublished data). The CAD5 PrP interactome is measurably different to other prion-susceptible cell lines^56^, and CAD5 cells release more exosomes and have lower basal autophagy levels than N2a cells^57^. The iterative single cell cloning process we undertook provides a series of related subclones with increasing susceptibility to prion infection across all assayed homogenates, rather than selecting for individual strains – this suggests enrichment of more broadly-relevant susceptibility factors. Determining sequential differences in gene sequence and expression in subclones with increasing prion susceptibility may help to elucidate these factors.

EKV cells and other humanised CAD5 subclones expressing valine at codon 129 (HuPrPV129) failed to propagate infectious prions after challenge with Type 2 MM sCJD homogenates. Tg152c mice expressing HuPrPV129 are susceptible to all human prion strains, regardless of genotype; in contrast, iPSC-derived astrocytes were only able to propagate sCJD prions in a genotype-dependent manner^43^. Whilst transmission of Type 2 MM sCJD prions to a HuPrPV129 host is possible in transgenic mice, the onset of clinical disease, and accordingly the prion replication rate, is much slower^58^; it is feasible that this is insufficient to sustain infection in culture. Interestingly, we separately developed CAD5 HuPrPM129 subclones, which to date do not clearly replicate PrP^Sc^ from Type 2 MM sCJD homogenates, but do propagate highly infectious Type 4 vCJD prions (manuscript in preparation). CAD5 cells, or our humanised subclones, may lack the appropriate cellular milieu to propagate Type 2 MM sCJD, or this may be a function of detection – MM sCJD strains are approximately ten-fold less PK-resistant, and have a greater proportion of PK-sensitive material^59^. Further screening of humanised HuPrPM/MV129 subclones, combined with orthogonal detection methods such as alternative protease digestion^30^ and seeding assays, may lead to a clone able to propagate Type 2 MM sCJD prions. Ultimately, it is likely that a panel of different cell lines will be required to replicate the entire gamut of human prion strains^7^.

At present, the EKV cell Human Prion Assay has a mean lower inoculum concentration detection limit of 10^-5^ across assayed brain homogenates. Limited data exist regarding the titre of human prions in various tissues or biofluids^60^, in large part due to the prohibitive resources required to perform these experiments in animal bioassays. Recent evidence points to higher infectivity than previously suspected in non-neural samples such as blood plasma^61^, skeletal muscle and exocrine glands^17^, raising valid concerns over transmission risk. A particularly appealing prospect is the capacity to quantify infectious human prions in non-neural tissues and biofluids, both to undertake a detailed survey to definitively address infection risk, but also to act as an *in vivo* biomarker of disease. The current detection limit is unlikely to support this, but several potential avenues to further increase cell susceptibility warrant further investigation^62, 63^.

## Materials and Methods

### Cell lines and routine culture

Unless otherwise specified, CAD5 cells and subclones were cultured in Gibco Opti-MEM™ Reduced Serum Medium containing 10% Bovine Growth Serum (ThermoFisher, HyClone™) and 1% penicillin/streptomycin (ThermoFisher, 10,000 U/ml). Cells were routinely passaged 1:6 – 1:10 every 3-5 days via gentle trituration, and maintained in a humidified incubator (5% CO2, 37°C).

### Retroviral expression and reconstitution of CAD-KDB3 cells with human PrP

Derivation of CAD5-KDB3 cells, in which the endogenous mouse PrP has been stably silenced by RNA interference, has been described in Bhamra *et al*^23^. To reconstitute KDB3 cells with human PrP, a retroviral expression construct was prepared by inserting the human PrP open reading frame with valine at codon 129 into pLNCX2. The human PrP open reading frame was synthesised (Invitrogen GeneArt, ThermoFisher). In addition to valine at codon 129, the human signal peptide sequence (aa 1-22) was replaced with the mouse signal peptide: mouse signal sequence MANLG<u>Y</u>W<u>L</u>L<u>A</u>LFV<u>TM</u>W<u>T</u>D<u>V</u>GLC (underlined amino acids indicate deviation between mouse and human amino acids). The human PrP coding region was sequence verified in two independent constructs (pLNCX2 mssHuPrPV128-8 and -24).

Both pLNCX2 constructs were packaged into recombinant retroviruses by transient transfection into Phoenix ecotropic cells (ATCC, LGC Standards, Middlesex, UK) in conjunction with vesicular stomatitis virus G (VSV-G) envelope expression vector (pMDG2) by FuGENE 6 Transfection reagent (Promega), according to the manufacturer’s instructions. We have found that pseudotyping the ecotropic retrovirus particles with the VSV-G protein increased the efficiency of stable transduction in CAD5 cells. Stable transduction of cells with pLNCX2 retroviruses was achieved by antibiotic selection using geneticin (G418, ThermoFisher, 400 µg/ml final concentration). 240 single cell clones were isolated for each construct and expanded.

### Human Prion Assays

Frontal cortex 10% weight/volume homogenates were prepared by homogenisation in sterile Dulbecco’s phosphate-buffered saline lacking Ca^2+^ and Mg^2+^ ions (D-PBS), using Duall tissue grinders. Initial human prion infectivity assays used the same format as the previously described Scrapie Cell Assay^11^, but with modifications to enable detection of infectivity of human prions. 18,000 cells were seeded in a 96-well microplate and incubated for 24 hours in 200 µl growth medium, after which 100 µl medium containing unfiltered prion-infected brain homogenate was added to each well. Cells were passaged twice weekly 1:6 / 1:8 at 3 - 4 day intervals, respectively. At confluence after passage 3, 4, and 5, 18,000 cells were filtered onto pre-activated ELISPOT plates (MultiScreenHTS IP Filter Plate, 0.45 µm, Merck Millipore), fixed at 50°C for 1 hour, then lysed and incubated at 37°C for 1 hour in the presence of proteinase K (2.25 µg/ml), followed by 80 µl phenylmethylsulfonyl fluoride (PMSF, Sigma) for 10 minutes at room temperature. 80 µl of a 1:25000 dilution of ≥ 250 units/µl stock benzonase solution (Merck Sigma) was then added and the plates incubated at 37°C for 15 minutes. PrP^Sc^ was detected using 0.5 µg/ml mouse anti-PrP monoclonal antibody ICSM18 (D-Gen Ltd, UK), followed by alkaline phosphatase conjugated anti-mouse IgG1 and conjugate substrate (Bio-Rad). After drying plates, PrP^Sc^-positive spots were quantified using either the Zeiss KS ELISPOT system or the BioSys Bioreader®-7000-F imaging system.

For the optimised Human Prion Assay (HPA) for EKV cells, 2500-5000 EKV cells in 150 µl medium were added directly to wells of a 96-well microplate containing 150 µl brain homogenate diluted in medium. Cells were passaged as above, and 20,000 filtered per well for ELISPOT revelation after passages 4 and 5.

### Single cell cloning

To derive a pool of single cell clones by limiting dilution, a CAD5-KDB3-derived parent cell line was resuscitated from cryopreservation and passaged twice in 2 µg/ml puromycin and 400 µg/ml G418. Cells were seeded in 10 cm dishes at a density of 150 cells/dish in 15-20 ml selection medium, and incubated for 10-12 days. Cultures were reviewed periodically, and single cell colony locations marked on the dish. After 10-12 days (approximately 5-7 doublings), growth medium was carefully changed and single cell colonies aspirated into individual wells of 96-well microplates. Wells were passaged twice at confluence, then expanded to triplicate plates; two replicate plates were cryopreserved and the third assessed for human prion susceptibility, as above.

### Developing cells persistently infected with human prions and pilot therapeutic assays

EKV cells were resuscitated, passaged twice in selection medium and challenged with 0.001% human prion-infected brain homogenate, followed by five passages to dilute out the inoculum. At confluence after passage 5, cells were seeded for single cell cloning as above, at a density of 1 cell per well in 300 µl medium, in two to four 96-well microplates. Cells were allowed to grow to confluence, after which the media was replaced and cells passaged twice and expanded into triplicate plates. Two duplicate plates of each set were cryopreserved, and the final plate subjected to ELISPOT revelation to detect cells containing PrP^Sc^. Clones with the highest PrP^Sc^-positive spot number (infected EKV cells, iEKV) were expanded and cryopreserved in doubly sealed aliquots, in a dedicated vapour-phase liquid nitrogen storage tank exclusively for human prion-infected cells.

For pilot therapeutic assays, iEKV cells were resuscitated and passaged twice in growth medium. At confluence, 10,000 cells were seeded in 150 µl plain medium and allowed to adhere for 2 hours. A dilution series of anti-PrP ICSM18 or BRIC222 (an IgG1 isotype control mouse monoclonal antibody) were added in 150 µl medium to appropriate wells. After 5 days of incubation, media was gently replaced and 7,000 cells per well filtered for ELISPOT or toxicity assay. Untreated iEKV cells, non-infected EKV cells, media-only wells and DMSO-treated iEKV cells were used as controls. BRIC222 (IBGRL, Bristol) was a kind gift from Dr. Azadeh Khalili-Shirazi. XTT toxicity assay (CyQuant XTT Cell Viability Assay, ThermoFisher Scientific) was undertaken as per the manufacturer’s instructions.

### Transmission studies and immunoblotting

Tg152c mice were bred and maintained as described previously^65^. Strict biosafety protocols were followed throughout. Inocula were prepared using disposable instruments for each inoculum, in a microbiological containment level 3 laboratory, and inoculations performed within a class 1 microbiological safety cabinet. Inocula were prepared in D-PBS, using Duall tissue grinders for human brain tissue, or ribolysed in a Precellys®24 tissue homogeniser. Mice were culled when clinical signs of scrapie sickness were detected, or at 600 days post-inoculation, as described previously^65^.

For inoculation with cell homogenates, 81F9 or KDB3 cells were challenged with sCJD brain homogenate, or growth media only, and passaged six times. At confluence, cells were pooled, resuspended in D-PBS at a concentration of 10 million cells per ml, ribolysed and frozen at -80°C, for animal inoculation.

For neuropathology and immunohistochemical analysis, transgenic mouse brains fixed in 10% buffered formal saline were immersed in 98% formic acid for 1 hour. Following further washing in 10% buffered formal saline, tissue samples were processed and embedded in paraffin wax. Serial sections of 4 μm nominal thickness were prepared. Deparaffinized sections were analysed for abnormal human PrP deposition on a Ventana Discovery XT automated IHC staining machine (Roche Tissue Diagnostics). In brief, sections were treated with cell conditioning solution (Discovery CC1; Roche Tissue Diagnostics) at 95°C for 30 minutes, followed by treatment with a low concentration of protease (Protease 3; Roche Tissue Diagnostics) for 16 minutes. Anti-PrP monoclonal antibody 3F4 (BioLegend) was used in conjunction with a biotinylated polyclonal rabbit anti-mouse immunoglobulin secondary antibody and Ventana proprietary detection reagents utilising 3,3′-diaminobenzidine tetrahydrochloride as the chromogen (DAB Map Detection Kit). Conventional methods on a Gemini AS Automated Slide Stainer (Thermo Fisher Scientific) were used for hematoxylin and eosin (H&E) staining. Positive controls for staining were used throughout. All slides were digitally scanned on a Hamamatsu NanoZoomer 360 instrument, and images captured from the Hamamatsu NZconnect image management software for the preparation of light microscopy images.

Samples were processed for immunoblot detection of proteinase K-resistant PrP as described previously^66^. Human tissue samples were incubated for 1 hour at 37°C in 50 µg/ml proteinase K, and transgenic mice or cell lysate samples in 100 µg/ml proteinase K. CAD5 cell lysates were prepared from frozen pellets (10 million cells), resuspended in 1 ml D-PBS, ribolysed and immediately processed for immunoblotting.

### Statistical Analysis, Ethics, and Data Availability and Funding

All statistical analyses were performed using Graphpad Prism 9. All data referenced in this work are available on reasonable request. Ethical approval was granted by the local ethics board, and NHS Research Ethics Committee for human sample collection.

## Acknowledgements

We thank all the individuals, their caregivers and families who kindly donated material used in this research. We thank Richard Newton for graphics, Azadeh Khalili-Shirazi and Stephanie Canning for provision of BRIC222 and technical support, Adam Wenborn and Jonathan Wadsworth for considerable intellectual and technical support, and Juan Ribes and Peter Klöhn for technical support and materials. Akin Nihat was supported by a UK Medical Research Council Clinical Research Training Fellowship (grant ID MR/P019862/1). This work was funded by the UK Medical Research Council and the National Institute for Health Research UCLH Biomedical Research Centre.

## Author Contributions

PSJ and JC conceived the work. PSJ, JC and SM supervised the work and secured funding. AN secured funding, undertook experiments and data analysis, drafted and revised the manuscript. PA, CS, MR, JL, SB, PSJ undertook experiments and data analysis. All authors contributed to manuscript revision and reviewed final content.

